# Human Slip Control: Investigating the Role of Hand Acceleration Modulation in Preventing Slips

**DOI:** 10.1101/2024.12.29.630651

**Authors:** Kiyanoush Nazari, Willow Mandil, Marco Santello, Amir Ghalamzan

## Abstract

Ensuring a stable grasp during manipulative movements is crucial for robotic applications. While grip force has been the primary means of slip control, our human study revealed that trajectory modulation is also an effective slip control policy during pick-and-place tasks. Motivated by these findings, we developed and compared a slip control policy based on trajectory modulation to one based on grip force control for robotic pick-and-place tasks. Our results show that trajectory modulation significantly outperforms grip force control in certain scenarios, highlighting its potential for slip control in robotics. Moreover, we demonstrate the importance of incorporating forward models in developing effective trajectory modulation slip control systems. Overall, our study provides insight into an alternative method for slip control and suggests that humans use trajectory modulation as an alternative to grip force control for slippage prevention. This insight helps design algorithm for improving robotic manipulation tasks to prevent slippage.

## 1. Introduction

Stable object manipulation without unwanted slippage is a fundamental challenge in both human motor control and robotic systems. A stable grip is essential for ensuring that objects can be securely manipulated, particularly in dynamic environments. Traditionally, research has focused on grip force modulation as the primary mechanism for preventing slip, where a safety margin is maintained by exerting more grip force than the minimum required to hold an object. However, emerging evidence suggests that the human nervous system may use more sophisticated strategies, incorporating both grip force control and motion trajectory modulation. This paper investigates the hypothesis that humans utilize a combination of these strategies to prevent slip during object manipulation tasks, offering a broader perspective on how the nervous system ensures stable object handling.

The classical understanding of slip control revolves around grip force modulation. Early studies, such as those by Johansson and colleagues, established that humans typically apply grip forces slightly above the minimum necessary to prevent slippage, creating a safety margin [1, 2]. This safety margin is adjusted dynamically through a combination of predictive and reactive mechanisms, incorporating both feedforward control and real-time sensory feedback [3]. As a result, grip force modulation has long been considered the cornerstone of human slip-prevention strategies.

Recent studies, however, have suggested that slip prevention may be a more complex process involving the integration of multiple control strategies. Previous work has shown that humans adjust their hand movements in response to environmental changes, such as the need to stabilize objects and prevent slippage when manipulating them [4, 5]. Building on this understanding, we hypothesize that grip force control is not the sole method employed by humans to avoid slips. In a human study involving a pick-and-place task, we discovered that increasing grip force is not always the primary strategy for slip control. Instead, participants modulated their hand velocity and acceleration as a means to stabilize the object and control slip. This finding challenges the prevailing view in the robotic community that grip force is the only effective method for slip control based on human studies [6]. Our results show that humans modulate their hand acceleration to efficiently stabilize objects and maintain a safe grip force, thus preventing slippage while avoiding excessive effort. These findings suggest that hand velocity modulation plays a significant role in slip control and highlight the need to explore both grip force and trajectory modulation in achieving stable object manipulation.

### 1.1. Background and Related Work

*Grip Force and Safety Margin* The concept of the ‘safety margin’ for slip prevention has been a central focus in the study of human motor control. Humans typically apply between 10% and 40% more grip force than necessary to prevent slip, compensating for various factors such as object weight, texture, and friction [2, 7, 8, 9]. This extra force is dynamically adjusted in response to sensory feedback, enabling fine-tuned grip force control during object manipulation. Johansson and colleagues demonstrated that the nervous system maintains grip forces slightly above the minimum required, adjusting the force based on the object’s properties and real-time feedback from the fingers [3].

The role of somatosensory feedback, particularly from mechanoreceptors in the skin, is crucial in regulating grip force and preventing slip. These mechanoreceptors provide information on texture, pressure, vibration, and skin stretch, enabling the nervous system to detect slippage and adjust grip force accordingly [10, 11, 12]. Feedback from both tactile and proprioceptive sensors helps the brain make rapid, reflexive adjustments to grip force, ensuring that objects are securely held during manipulation.

Reflexive adjustments in response to tactile input depend on the posture and positioning of the arm, indicating a close coupling between proprioception and tactile feedback [13]. These afferents respond to different phases of slippage, from the ‘stuck phase’ to ‘partial slip’ and ‘full slip’ [14, 15, 16]. The tactile feedback gathered from the fingers and hands is crucial for detecting slippage, enabling the brain to initiate appropriate motor responses to maintain object stability. In addition to tactile feedback, proprioceptive signals from muscles and joints also play a significant role in regulating grip force, further enhancing the ability to control object stability. The integration of these sensory signals allows for the precise modulation of grip force, optimizing it based on environmental conditions and object properties.

The neural basis for grip force regulation involves a distributed network of brain regions, with the basal ganglia playing a pivotal role in precision grip force control [17].

The nervous system integrates proprioceptive and tactile feedback in a tightly coordinated manner. It has been observed that caudal regions of the primary motor cortex (M1), which primarily govern hand movements, receive a higher proportion of tactile input compared to the rostral regions that control more proximal limb movements [18, 19].

Grip force regulation is governed by a hierarchical neural network involving multiple brain regions. Early responses to perturbations, such as the ‘short-latency reflex’ (SLR), are generated by spinal circuits and occur within 25–50 ms of a slip event [20, 21]. These quick reflexive actions help stabilize the object before voluntary control takes over. The ‘long-latency reflex’ (LLR), which occurs 50–70 ms after a slip event, involves both spinal and supraspinal pathways, including the primary motor cortex (M1) [22]. Voluntary responses, which follow after 100 ms, are modulated by a network of cortical and subcortical regions, including the premotor cortex and basal ganglia, allowing for more deliberate adjustments to grip force [23, 24]. These voluntary reactions are essential for fine motor control and are central to skilled object manipulation.

Despite extensive research on grip force control, several key gaps remain in our understanding of how humans prevent slip during object manipulation. First, the role of hand trajectory modulation in slip prevention has been largely overlooked, with most studies focusing exclusively on grip force adjustments. Second, the interaction between visual feedback and hand motion control, particularly in terms of maintaining object stability, is not fully understood. Finally, the relative contributions of predictive and reactive control mechanisms in trajectory-based slip prevention strategies have not been systematically explored. These limitations suggest that the current understanding of slip prevention is incomplete, and more research is needed to identify other potentially more efficient strategies.

### 1.2. Research Objectives and Contributions

This paper makes several significant contributions to the field of human motor control and slip prevention: (I) We present novel experimental evidence demonstrating that humans employ hand acceleration modulation as an effective strategy for slip prevention, challenging the traditional grip-force-centric view of object manipulation. (II) We quantify the relative contributions of grip force and hand trajectory modulation in slip prevention through carefully designed pick-and-place tasks under two distinct conditions, highlighting the importance of both strategies. (III) We propose a new theoretical framework that integrates grip force control and hand trajectory modulation, offering a comprehensive perspective on how the nervous system achieves stable object manipulation.

Our study provides a fresh perspective on the complex neural control mechanisms involved in slip prevention and offers valuable insights for future research on the integration of these control strategies in both human motor control and robotic systems. This result is significant because it demonstrates that humans modulate hand velocity/acceleration to efficiently keep the required grip force for slip prevention bounded to their affordable grip force [25].

### 1.3. Paper Structure

The remainder of this paper is organized as follows: Section 2 presents the research question and hypothesis, followed by a detailed human subject study that investigates slip prevention strategies. Section 3 outlines the experimental methodology and data analysis techniques used in our study. Section 4 presents the results, followed by an in-depth discussion in Section 5. Finally, Section 6 concludes the paper, highlighting future research directions and the implications of our findings for robotic systems.

## 2. Research Question, Hypothesis, and Human Subject Study

### 2.1. Do humans use hand movement modulation in addition to grip force control for preventing slippage?

Slip control during object manipulation is a crucial component of human motor behaviour, especially when interacting with objects that may slide or shift due to insufficient grip. Traditional theories of slip prevention have predominantly focused on grip force modulation, where humans apply grip forces above the minimum required to prevent slippage. This approach is grounded in the concept of a safety margin, which has been extensively studied and is regarded as a key mechanism for stabilizing objects during manipulation. However, recent studies suggest that grip force modulation alone may not be the sole strategy employed by humans to prevent slip. Rather, slip prevention is likely governed by a more sophisticated and dynamic integration of both grip force control and motion trajectory modulation. This section reviews the role of grip force control in preventing slippage, presents evidence that additional strategies may be at play, and highlights the importance of trajectory modulation in enhancing stability during object manipulation.

The coordination of tactile, proprioceptive, and visual feedback is essential for effective slip prevention during object manipulation. As objects are handled, the human nervous system continuously adjusts grip force to ensure that it exceeds the threshold required to prevent slippage. These adjustments are informed by sensory feedback that signals when a slip is imminent, such as changes in skin deformation or vibrations from the object [26].

In real-world conditions, the nervous system anticipates potential slippage by proactively adjusting grip force before it becomes necessary. This anticipatory mechanism creates a ‘safety margin’ that accounts for dynamic factors such as object mass, friction, and changes in orientation [27, 28]. The safety margin thus helps to prevent slippage by ensuring that the applied grip force remains above the critical threshold while avoiding unnecessary increases that could lead to fatigue or discomfort. This proactive adjustment enables the human motor system to balance grip stability and energy efficiency during object manipulation.

Research also shows that grip force is not solely determined by the object’s weight or surface properties. Instead, it is dynamically adjusted based on continuous sensory input, ensuring that the applied forces remain within an optimal range that prevents slippage without overexerting effort [2, 29]. This integration of sensory feedback and motor control is a robust mechanism that enables slip prevention even in dynamic or uncertain environments, where conditions constantly change and new challenges arise [30].

Despite the well-established role of grip force modulation in slip prevention, recent studies suggest that the interaction of trajectory modulation with grip force control—especially during complex tasks—has yet to be fully explored. Human manipulation is often characterized by both time-sensitive goals (e.g., completing a task quickly) and stability requirements (e.g., maintaining control over the object). The control of hand trajectory in robotics, including velocity and acceleration, can help maintain object stability, particularly in scenarios where the forces acting on the object are not constant [31, 32]. In such cases, humans may adjust their hand motion to stabilize the object and prevent slippage, rather than relying solely on increasing grip force.

Our hypothesis builds on this understanding: while grip force modulation remains an essential strategy, hand trajectory modulation also plays a significant role in preventing slip. Recent findings suggest that humans utilize hand velocity and acceleration adjustments as a means to stabilize objects during manipulation tasks, particularly when direct slip control via grip force is insufficient or when environmental conditions demand fine-tuned adjustments to object handling [4, 5]. This dual strategy—integrating both grip force control and trajectory modulation—may provide a more flexible and efficient means of maintaining stability, especially in tasks that require both stability and speed.

Despite the importance of grip force control in slip prevention, several key gaps remain in our understanding of the mechanisms involved. First, the role of hand trajectory modulation in slip prevention has been largely overlooked, with most studies focusing exclusively on grip force adjustments. Second, the interaction between visual feedback and hand motion control, particularly in terms of maintaining object stability during manipulation, is not well understood. Finally, the relative contributions of predictive (e.g., trajectory modulation) and reactive (e.g., grip force adjustments) mechanisms in slip prevention strategies have not been systematically explored. These gaps suggest that current models of slip prevention may be incomplete and that a more integrative approach is needed to understand the dynamics of object handling fully.

### 2.2. Hypothesis and human study

#### Hypothesis: Human Hand Motion Modulation for Slip Control. Participants

To ensure the efficacy and validity of our human participant tests, we conducted over 6 preliminary experiments in a pilot study to optimise our process and scores. Sixteen right-handed participants (eight male and eight female), aged between 20-40, provided informed consent and participated in the tests. Ethics approval was obtained (Ethics Reference: UoL2022 10370) to conduct the experiments with human participants.

#### Experimental Setup

Each participant wore a glove equipped with a 9-axis inertial measurement unit (IMU, adafruit BNO055) and stood in front of a table (70×150×70 cm) with their feet positioned in a designated area at the centre of the table (See the figure in Table 1). They grasped a book (24×16×2cm, 300g) with the tips of their right hand’s thumb and middle finger. The middle finger was preferred over the index finger as it can generate greater flexion force [33] and made keeping the object in an upright pose easier for the participants. To ensure consistent grasping location across participants, the thumb was placed on a mounted tactile sensor (uSkin, XELA ROBOTICS), and the middle finger was placed on an attached part on the opposite side of the book (mirrored from the tactile sensor). The tactile sensor measured 3-axis calibrated force readings for 16 sensing elements (taxels) arranged in a 4×4 configuration within a 2.3×2.3 cm area. The table had designated zones for the start and end parts of the motion. Two visual markers (ArUco) were used to measure the task start and end time. A RealSense d435i camera measured the 3D pose of the markers. The data reading frequency for the tactile sensor, IMUs, and marker pose were 100, 60, and 60 Hz, respectively. We used the *ApproximateTime policy* from ROS to synchronise the sensory data. The first and second IMUs measured the acceleration and absolute orientation of the participant’s hand and the object, respectively.

**Table 1:**
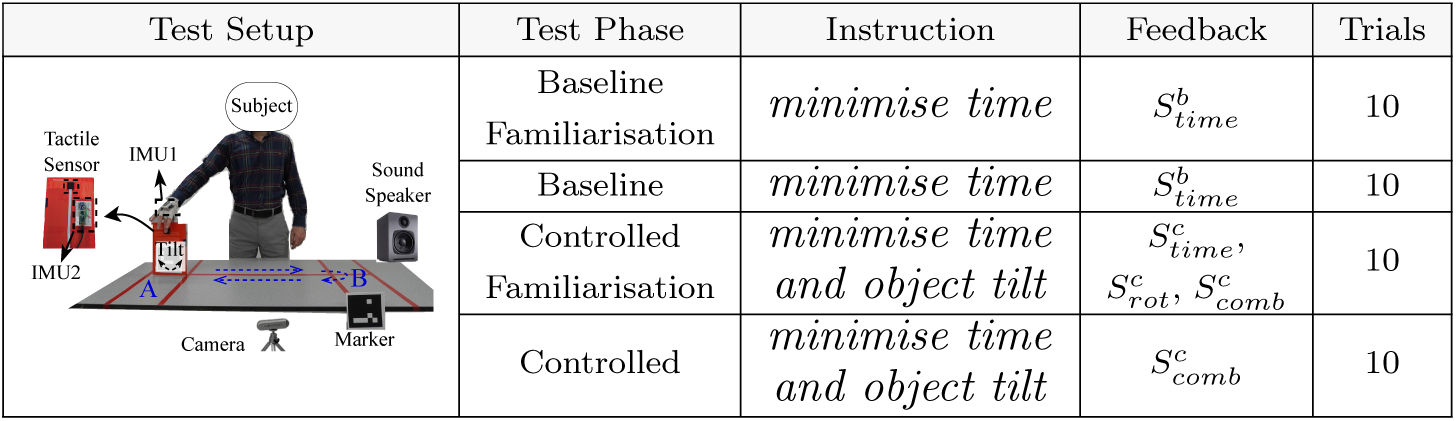
Human participant test for slip control analysis. IMUs measure participants’ hand acceleration and the object’s in-hand rotation angle around the grip axis (tilt). Audio feedback with *”beep”* sounds informs the participants when to lift and start the transport. The Test Phase column is sorted based on the chronological order of undergoing the tests. The superscript *”b”* and *”c”* for the feedback scores refer to B-Tests and C-Tests, respectively.

## 3. Experimental methodology and data analysis

The experimental procedure consisted of the following steps. First, the participants lift the book and perform a round move following the straight line between the start and end zones, as shown in the figure in Table 1. The test consisted of two parts, namely, *(i)* baseline and *(ii)* controlled. Each part included 20 trials, where the first 10 trials were referred to as *‘familiarisation trials’*, during which the participants learnt about the scoring system. The remaining 10 trials were referred to as *‘test trials’*. Test scores hyperparameters and tests conditions are designed based on a pilot study with 12 participants conducted prior to the real test.

The baseline task aimed to observe the participants’ hand motion profiles when they focused solely on minimising task execution time. To ensure consistency in task execution, we provided participants with a time score after each trial and encouraged them to try to improve their score in subsequent trials. The time score was calculated by discretising the sigmoidal shape function in equation (1) applied to the time-to-completion for all the participants as shown in Fig. 3. The score values were colour-coded to encourage higher scores, with the values *”Slow,” ”Normal,” ”Fast,”* and *”Excellent”* represented by red, yellow, light green, and dark green, respectively.

Since qualitative terms such as *”fast”* or *”minimum time”* can be perceived differently by different individuals, we show the score number in addition to semantic feedback to reinforce the participants to optimise the score. This provides participants with a more objective measure for the time-tocompletion.

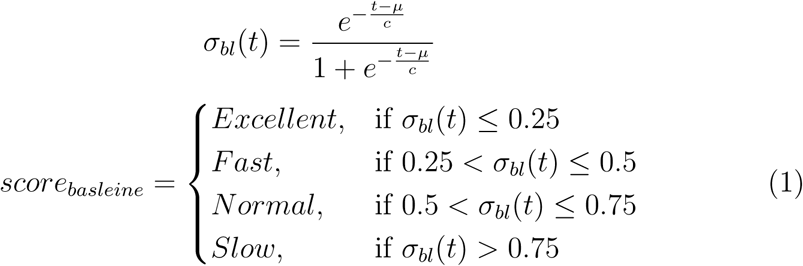

The shape and parameters of the sigmoid function in equation (1) (mean *µ* = 0.95 and slope *c* = 0.3) are determined based on the pilot test study.

In the controlled condition, we instructed the participants to perform the same moving task in the minimum time possible, while also aiming to minimise a second constraint: the rotation of the object around the grasp axis. Since participants were instructed to keep their hand pronated and the maximum hand rotation was 2.61% of the absolute object rotation (≡ 1.91*^◦^*) across all trials, controlling the object rotation is equivalent to controlling the rotational slip of the object around the grip axis. During the familiarisation trials, we evaluated the participants based on three scores: (1) *time score*, (2) *rotation-score*, and (3) *combined-score* for each trial. The time score is indicative of the time-to-completion, the rotation-score measures the angle of rotation around the grasp axis, and the combined-score is the weighted sum of these two scores. Each score is calculated as follows:

### Time score

we calculate the participant’s time score in the C-tests by normalising the time-to-completion with the average time-to-completion across the 10 B-tests. This represents how fast the participant performs the task compared to the B-tests. This normalised time reduces the variation among participants compared to the raw time-to-completion in our pilot tests. To formalise this, let *t_µ_* denote the average time-to-completion in the B-tests and *t_i_* denote the sample time-to-completion in the FC-tests and C-tests. The normalised time is calculated as 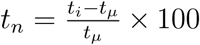 We calculate the continuous time score *S_t_* using the sigmoid function *σ*(*t*) in equation (2), which

is shown in Fig. 2(a).

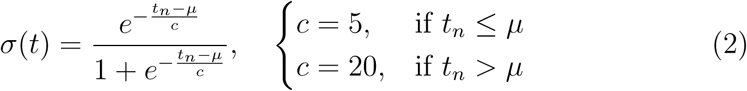

Where *µ* = 60. The function has an asymmetric shape to give more weight to slower times, as deviations from the optimal time are more detrimental than faster times. The y-axis of Fig. 2(a) illustrates how *S_t_*is discretised into the categories of *Slow*, *Normal*, *Fast*, and *Excellent* using colour-coded values of red, yellow, light green, and dark green, respectively.

**Figure 1:**
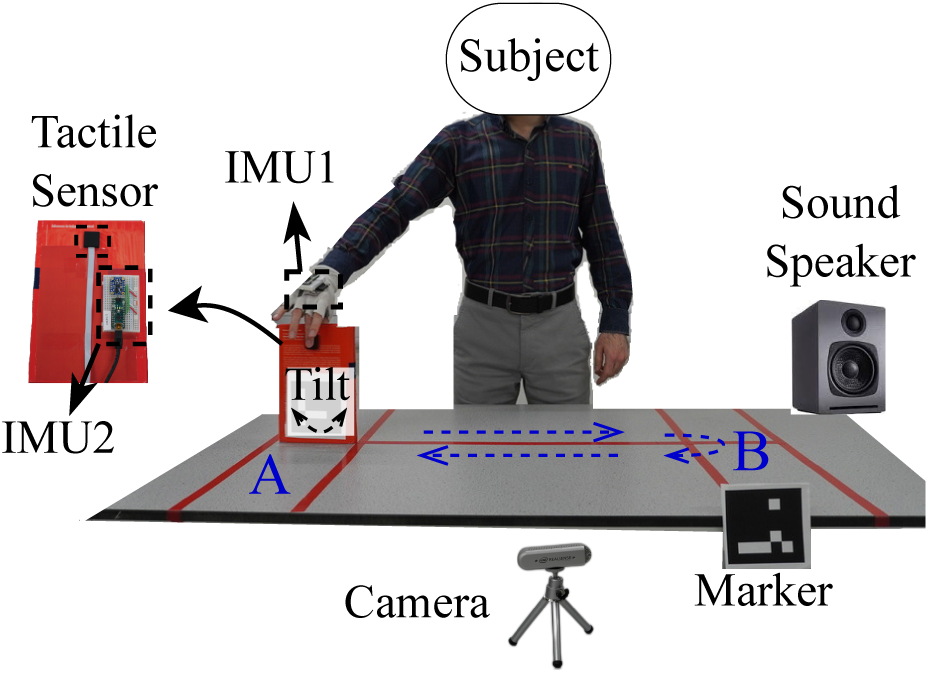
Human physiology test set up. Inertial Measurement Units (IMUs) and visual markers help measuring participants hand acceleration, task completion time, and object rotation during task execution. After every five trials, participants had two minutes resting time to avoid high muscle fatigue.

**Figure 2:**
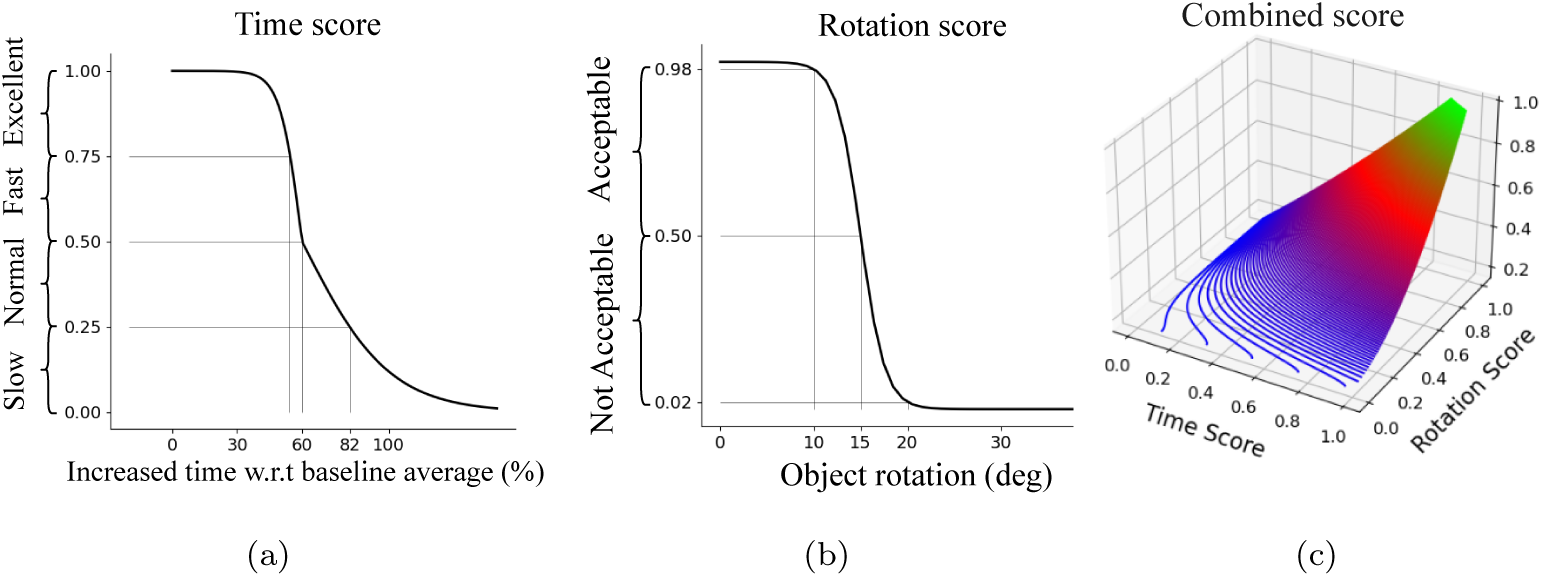
Participant scores for the controlled task. (a) time score, (b) rotation-score, and (c) combined-score.

**Figure 3:**
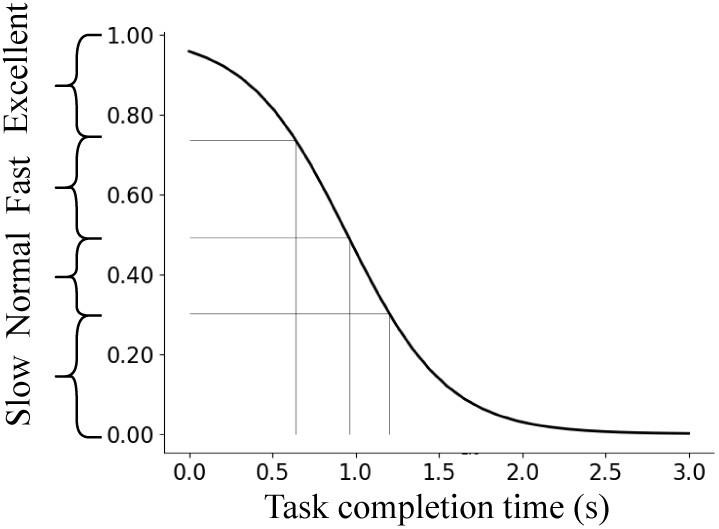
Baseline time score.

### rotation-score

To evaluate the participants’ ability to control rotational slip, we use an IMU to measure the rotation of the grasped object around the grip axis. We apply a sigmoid function, given by 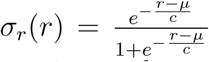 calculate the rotation-score. Where *r* denotes object rotation and *µ* = 15 and *c* = 1.3 are the parameters of the sigmoid function, chosen based on the pilot study. The rotation-score *S_r_* is derived by discrtising *σ_r_*(*r*). In the controlled condition, the participants are instructed to minimise object rotation while completing the task in minimum time. To provide feedback to the participants, we colour code the rotation-score to indicate if the participant’s performance was *acceptable* (green) or *not acceptable* (red). This encourages the participants to balance their focus on both rotation and time. The sigmoid function for rotation-score calculation helps us to map the continuous rotation value to a score between 0 and 1. The sigmoid function has a symmetrical shape, as shown in Fig. 2(b), and its discrete version shows the acceptability of the participants’ performance.

### combined-score

In a pilot study where we presented the participants with only their time and rotation-scores, we observed that many participants focused on maximising only one of the scores in each trial, rather than trying to improve both scores simultaneously. To encourage the participants to balance their focus on both time and rotation, we designed a third score that combines the effects of the time and rotation-scores.

The combined-score is calculated by the function in equation (3) using the normalised time score (*S_t_*) and the rotation-score (*S_r_*):

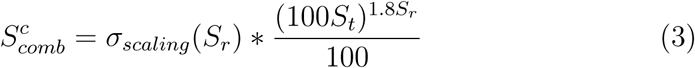

Fig. 2(c) shows the combined-score function which is designed heuristically based on a series of pilot studies. The resulting score reflects how well the participant performed the task with respect to both time and rotation-scores. We use the combined-score in the C-tests to encourage participants to balance their focus on both time and rotation. In the FC-tests, we provide all three scores to the participants to help them become familiar with the performance metrics. However, we only provide the participants with the combined-score during the C-tests. Participants performed 10 FC-tests and 10 C-tests.

Separate one-way repeated measure analysis of variance (ANOVA) was performed using SPSS Statistics with Trial (10 levels) and Condition (2 levels; baseline and controlled) as within-subject factors for each feedback score. Multivariate analysis of variance (MANOVA) was performed jointly on participants task time-to-completion, object maximum rotation and maximum grip force in the FC-tests and C-tests to find statistically significant changes of these variables. The threshold for statistical significance is chosen as *p* = 0.05 for both ANOVA and MANOVA tests. Separate Spearman’s Rank-Order Correlation were performed using SPSS Statistics to analyse the correlation between participants’ task time-to-completion, maximum object rotation, and maximum grip force.

## 4. Results

In this study, we took inspiration from the human control policy for slip control. We compare the B-Tests to a robot executing a pick-and-place task following a pre-defined reference trajectory without considering the tactile readings for controlling the movements. On the other hand, the C-Tests (namely, closed-loop system) use a learnt forward model to adjust the reference motion to minimise the likelihood of slips.

We present the results of sixteen participants lifting a book sitting on a table and moving it sideways from right to left for 67 cm and moving it back to the original point (the coordinates of the start and return points on the table are A=(33, 33) and B=(100, 33) cm, respectively, with respect to the table’s right-bottom corner shown in the figure of Table 1. Prior to the B-test and C-test, participants underwent 10 familiarisation (**F**) tests for each condition. In total, each participant performed 40 tests (i.e., 10 repetitions for Familiarisation B-Tests (*FB-tests*), *B-test*, Familiarisation CTests (*FC-tests*) and *C-test*). An entire test took less than an hour for each participant.

Each trial was followed by performance feedback, consisting of a **baseline**
time score 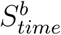 and a **controlled time score** 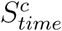, **rotation-score** 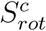, *comb* and **combined-score** 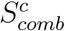 Table 1 shows the corresponding score *comb* feedback for each test phase. 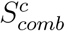 combines the impact of time and rotation in a single score for the C-Tests condition. All scores were normalised on a scale of 0 to 1, with higher values indicating better performance. We captured data of participants’ hand motion profiles using wearable Inertial Measurement Units (IMUs), object rotation with a second IMU attached to the object, and grip force measured by a 4× 4 uSkin tactile sensor [31, 34, 35] located under the participants’ thumb. We also used visual markers and the camera to measure time-to-completion for each participant. ANOVA and Spearman’s correlation analysis were used to test various experimental effects. A *p* value of 0.05 is considered statistically significant.

Fig. 4 illustrates the performance scores of all sixteen participants in the four test sections. We observe a steady increase of the time score in FB-test (Fig. 4(a)) showing participants’ learning curve, whereas reaching a plateau of the time score in B-test (Fig. 4(b)). Repeated measures ANOVA indicates that participants’ performance significantly improves during the FB-tests (*p <* 0.001), and remains stable during subsequent B-tests where no statistically significant changes of the score were found by ANOVA with *p* = 0.56. In the initial stages of the FC-tests (Fig. 4(c) and (e)), the participants rely on the model learnt in B-tests and prioritise the minimum time-to-completion objective, leading to a higher time score but a lower combinedscore. However, they quickly learn to relate their movements to the new scores in FC-tests and realise that the movement control policy learnt in FB-tests and used in B-test is not effective for maximising the rotation-score and combined-scores. Hence, they find an improved control policy in FC-tests to maximise the combined-score in Fig. 4(e) and use it in C-tests. Fig. 4(f) shows that during C-tests, the participants maximise the combined-score using the learnt model from FC-tests and improve their scores over time using the feedback score. Note that in C-tests, participants only have access to the combined-score. However, during FC-tests, they learnt the dynamics of the combined-score by receiving all three scores.

**Figure 4:**
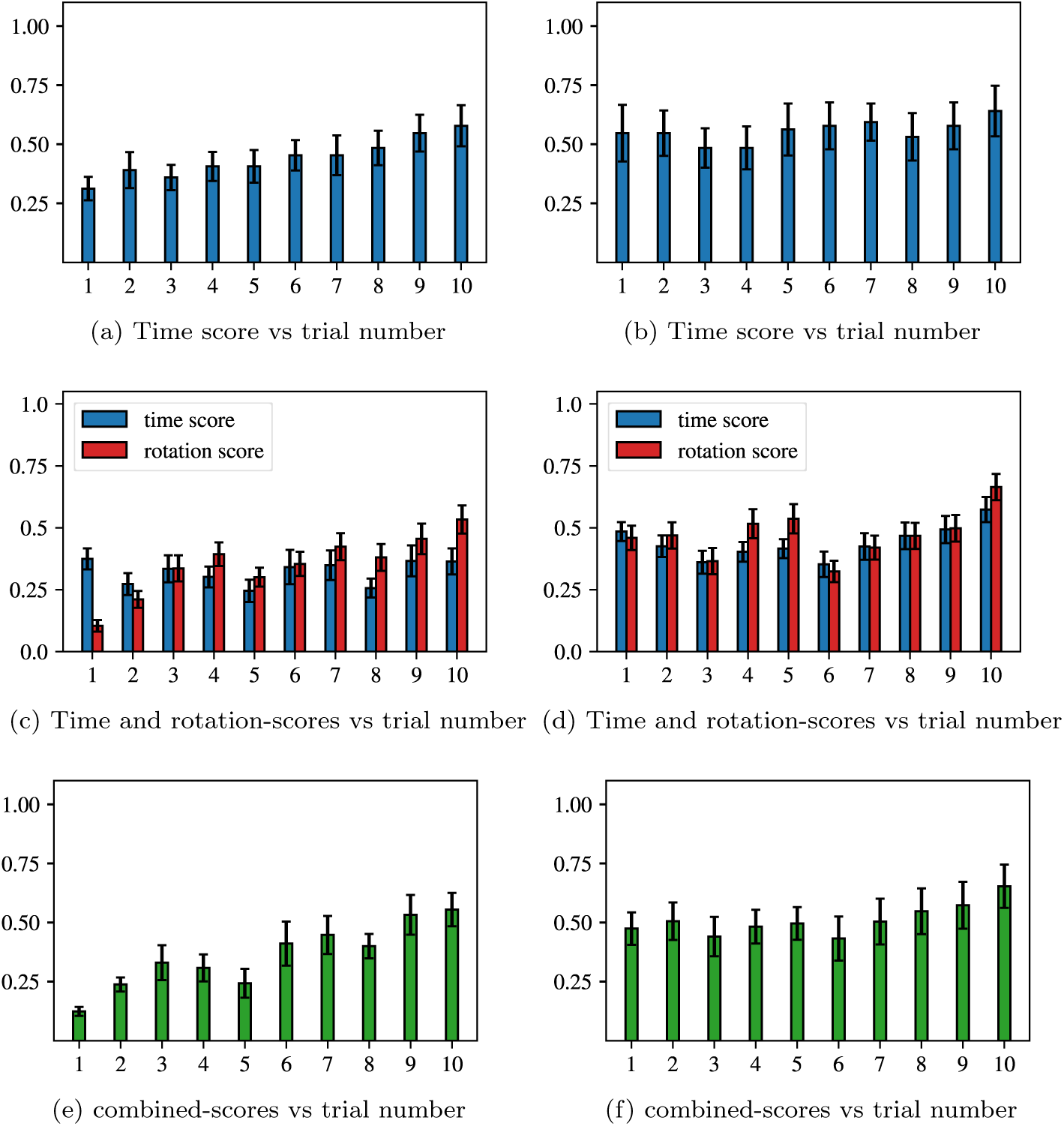
The mean and standard error of the mean of the performance scores for all sixteen human participants in slip control via trajectory modulation tests. *Familiarisation* tests (a, c and e): (a) time score in Baseline Tests, (c) time and rotation scores in C-Tests, and (e) combined score in C-Tests. Main tests (b, d and f): (b) time score in B-tests, (d) time and rotation-scores in C-Tests and (f) combined-score in C-Tests.

A pilot study that compared the performance of participants with having either all three or just the combined score in C-tests revealed that the latter strategy enabled the participants to more efficiently maximise the combined score. Repeated measures ANOVA indicates a significant increase of combined-score during FC-tests (*p* = 0.003) showing participants’ learning curve and no statistically meaningful change of combined-score in C-tests (*p* = 0.45). A one-way repeated measures MANOVA test showed a statistically significant increase in task time-to-completion (*p <* 0.001) and reduction in object rotation (*p <* 0.001) in C-tests compared to B-tests, proving that participants adapt their control policies based on performance feedback. This observation indicates that the participants sacrificed the minimum timeto-completion objective which is equivalent to reducing hand acceleration during C-tests to be able to control the object pose. Participants were instructed to keep their hand pronated and controlling the object rotation is equivalent to controlling the rotational slip of the object around the grip axis. The feedback in FC-tests and C-tests (namely rotation-score and combinedscore) involves the online acquisition of tactile feedback as well as visual feedback, even though the latter operates at longer sensorimotor delays than touch.

### 4.1. Human Grip Force Analysis

One possible explanation for the reduced object rotation in the C-tests is that participants used larger grip force relative to the B-tests. Fig. 5(a) displays the grip force exerted by a participant on the object during the Btests and C-tests. We, unexpectedly, observe decreased grip forces in the C-tests compared to the B-tests. Fig. 5(b) shows the maximum grip force during the motion execution, indicating a reduction in grip force values in the C-tests compared to the B-tests. Towards the end of the ten trials, the optimal grip force settled between 20 N to 22 N and between 14 N and 15 N for B-tests and C-tests, respectively. A repeated measures ANOVA for revealing the effect of task type on participants’ maximum grip force showed that the maximum grip force used during C-tests was close to being significantly smaller than during B-tests (P = 0.06). It has been reported that participants exert grip force slightly above the minimum grip force required to prevent object slip, and this ‘extra’ force has been defined as ‘safety margin’ [1]. The close to significant statistical changes could be due to the fact that the grip safety margins can vary widely across participants [36, 2, 37, 38] and there is large variability across subjects for maximum grip force ranging from 17.37 to 68.28 N in B-tests.

**Figure 5:**
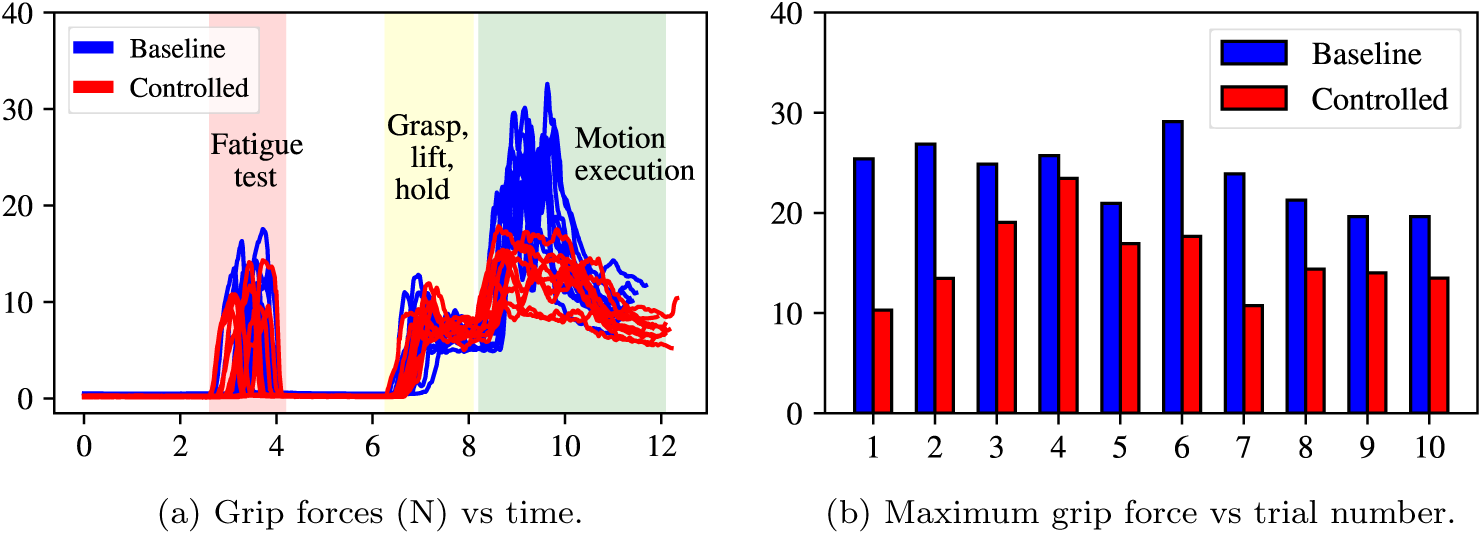
Grip force of one representative participant in Baseline and C-Tests. (a) Continuous values of participant’s grip force in Baseline (blue line) and C-Tests (red line) trials. The grip force is calculated by summing the normal force values for sixteen uSkin sensor taxels. Before each trial, the participants apply a firm grip and release to examine their physical fatigue as a confounding factor in the study. (b) Participant’s maximum grip force in the trials shown in (a).

The reduction in grip force could be attributed to fatigue. Hence, to assess the potential impact of physical fatigue on grip force, we conducted a ”fatigue test” before each trial where participants exerted a firm grip and release on the uSkin sensor, as shown in Fig. 5(a). We used the exerted normal forces over the 40 trials to evaluate the level of hand fatigue. To account for the potential confounding effect of physical fatigue, we normalised the grip force by the corresponding maximum normal force during the fatigue test press. As shown in Fig. 6(a), the resulting maximum normalised grip forces (average of sixteen participants) still showed a lower maximum grip force in the C-tests compared to the B-tests after considering the effect of physical fatigue. A repeated measure ANOVA test with *p* = 0.002 shows a significant reduction of maximum normalised grip forces in C-tests compared to B-tests. This indicates that the smaller grip forces observed in the C-tests were not solely due to physical fatigue, supporting our hypothesis that grip forces were modulated in response to the task requirements. The participants learnt that hand acceleration modulation is a less effortful approach for reaching grip safety margin than purely modulating the grip force in this case. Additionally, we analysed the maximum grip force during the fatigue test phase across all participants and presented the mean and standard error of the mean in Fig. 6(b). Repeated measures ANOVA shows that there is no specific or steady pattern of decrease in participants’ ability to grip (*p* = 0.23) within the 40 trials, even though the grip force values in C-tests are less than the B-tests. These findings rule out the potential confound of physical fatigue in our experimental results. The analysis of participants’ grip force addresses the first part of our first hypothesis, stating that humans do not solely rely on increasing grip force for object stabilisation in certain manipulation scenarios.

**Figure 6:**
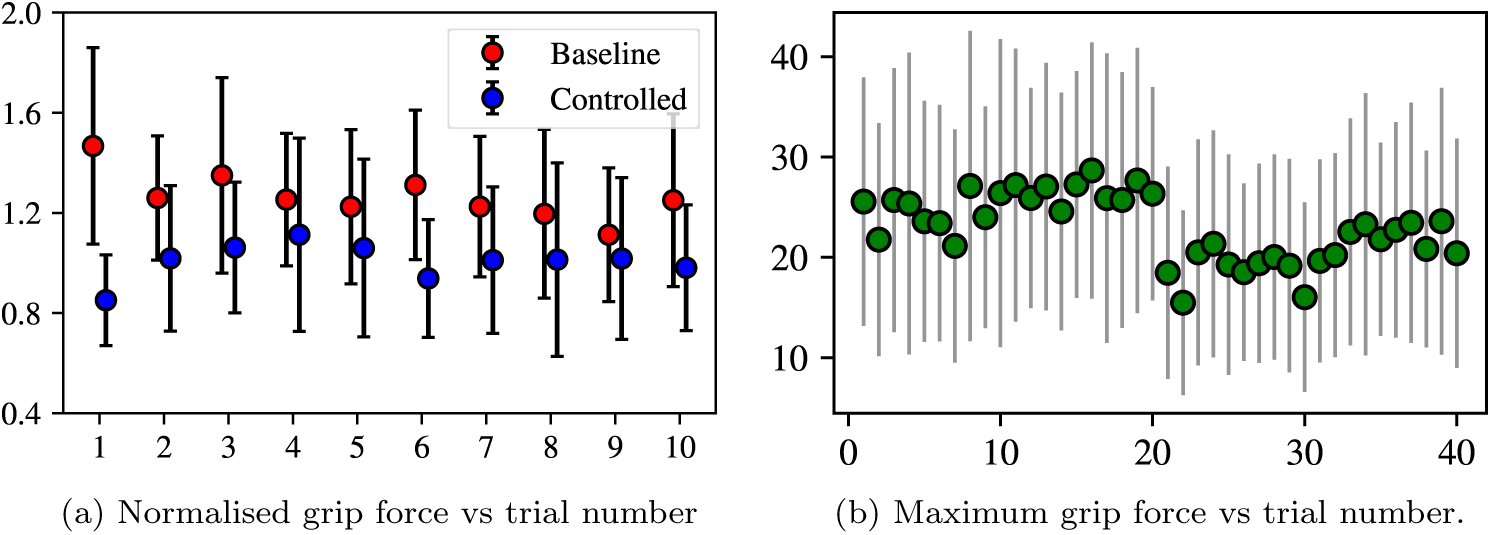
Analysis of fatigue confounding factor. (a) The maximum value of normalised grip force in Baseline and C-Tests. The raw grip force in the motion execution phase (see Fig. 5(a)) is normalised w.r.t the maximum value of the grip force in the fatigue test phase. The maximum grip force in the fatigue test phase for all test trials (n=40). The mean and standard error of the mean in both (a) and (b) are shown for sixteen participants.

An alternative policy to achieve the same performance scores (i.e object rotation) in the C-tests would be to purely increase the grip force rather than modulating hand acceleration. As a way to quantify how much grip forces were required to induce the same object rotations in C-tests when moving with hand acceleration trajectories used in B-tests, we simulated the dynamic equations of motion. Fig. 7 shows the exerted force on the book when transported in our manipulation task. We assume a circular contact zone of radius *R* between the finger pads and book cover with a friction coefficient of *µ*.

**Figure 7:**
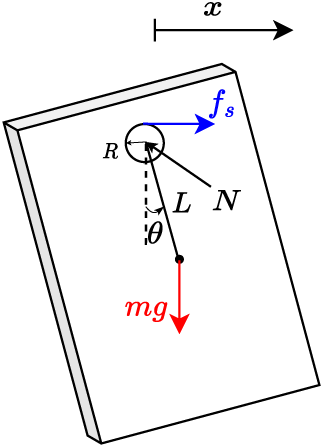
Grip force simulation.

The two degrees of freedom of the book consist of the translation *x* along the instructed path and rotation *θ* around the grip axis. Writing the dynamic equations for the rotational degree of freedom *θ* reads:

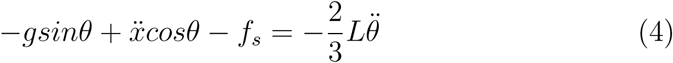

The objective is to find the frictional force *f_s_* when object rotation *θ* is coming from the C-tests and hand acceleration *x*ẍ is coming from the B-tests. Then by having *f_s_* = *µ* × *N* we can retrieve the normal grip force *N* . Fig. 8 shows the sum of the square root of the second norm of the grip force *N* in the 10 trials of B-tests, C-tests, and simulated tests.

**Figure 8:**
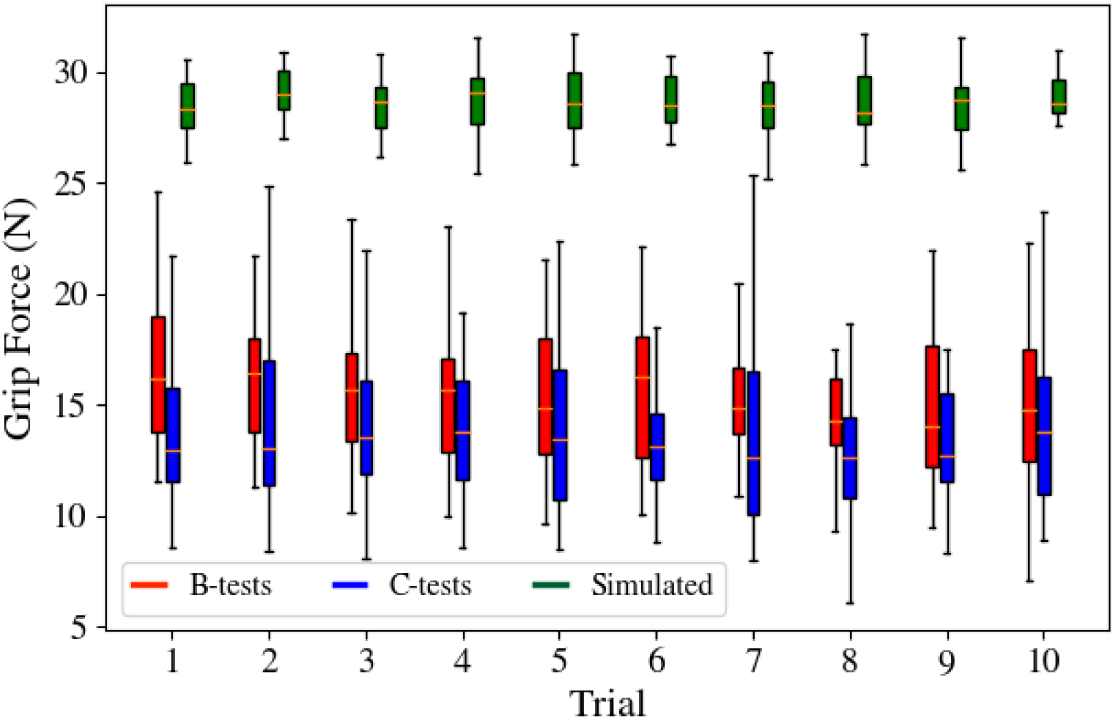
Grip force simulation results. Average grip force for 16 participants in 10 trials of the B-tests and C-tests is shown.

A one-way repeated measures ANOVA indicated the simulated grip force values were significantly larger than the grip force used B-tests and C-tests (*p <* 0.001). On average, simulated grip force (M = 28.58, SD = 0.17) was higher than B-tests (M = 15.58, SD = 0.59), and C-tests (M = 14.18, SD = 0.35). Post hoc comparisons were conducted using Bonferroni corrections. The difference between the average grip forces in B-tests and C-tests, 1.40 95% CI [0.87, 1.93], was statistically significant (*p <* 0.001). The difference between C-tests and simulated grip force, -14.40 95% CI [-14.63,-14.16], was statistically significant (*p <* 0.001). The difference between B-tests and simulated grip forces, -12.99 95% CI [-13.45, -12.54], was also statistically

significant (*p <* 0.001). The difference between simulated and actual grip forces (B-tests and C-tests) was much larger than the difference in grip forces used in B-tests versus C-tests

This indicates that subjects spontaneously chose to modulate their hand acceleration as a more efficient/feasible strategy for reducing the object rotation rather than keeping the same speed with increasing the grip force by 75%.

### 4.2. Hand Trajectory Modulation

To assess the impact of learning on hand acceleration profiles, we analysed the acceleration profiles of the participants in B-tests and C-tests (Fig.9(a) and (b), respectively). In the B-tests, the participant’s hand acceleration profiles were smooth, indicating that they learnt a motion trajectory and repeated it without making real-time modifications (i.e. an open loop control w.r.t sense of touch). A repeated measure ANOVA test with *p <* 0.001 shows a significant reduction of maximum hand acceleration in C-tests compared to B-tests for all participants. This control policy resulted in a range of object rotations between -76.81*^◦^* and 33.90*^◦^* (Fig.9(c)). In contrast, the hand acceleration profiles in the C-tests (Fig.9(b)) were characterised by high-frequency oscillations, indicating that participants were adapting their movements in real-time based on the sensory inputs (namely visual and tactile feedback). Given that humans generally respond faster to tactile cues than visual cues [39], tactile feedback may have played a crucial role in controlling the action. The high-frequency nature of the oscillations in hand acceleration profiles in C-tests supports this hypothesis that participants relied primarily on tactile feedback to close the control loop and prevent large rotation values. Consequently, the range of object rotation values in this condition (Fig. 9(d)) remained within a narrower range compared to the baseline (*p <* 0.001). Spearman’s correlation tests show a significant negative linear correlation between time-to-completion and maximum object rotation with *ρ* = −0.91. These findings suggest that learning the task allowed participants to adapt their control policies based on real-time sensory feedback and emphasises the importance of tactile feedback for trajectory modulation control policy in slip control.

**Figure 9:**
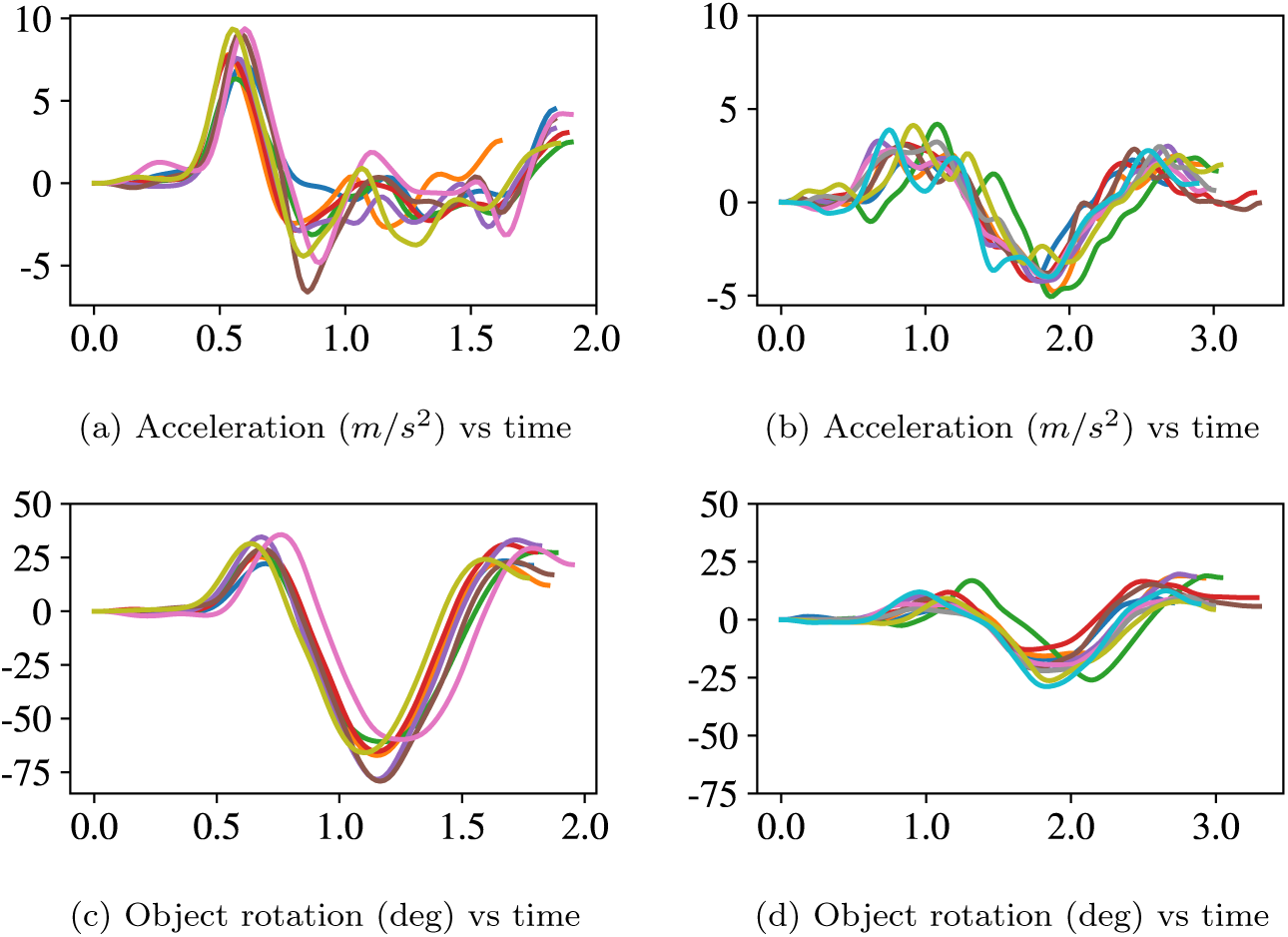
Hand acceleration and object rotation in B-tests and C-tests for a sample participant. (a) Hand acceleration in B-tests, (b) hand acceleration in C-Tests, (c) object rotation in B-tests, and (d) object rotation in C-Tests.

Based on the data presented in Fig. 5 and 9, it is evident that participants minimised the risk of object slip by reducing object rotation, which in turn was accomplished by employing a control policy in C-tests resulting in a reduced magnitude of the hand maximum acceleration. The modulated hand acceleration with high-frequency oscillations (which is a feature of feedback control using tactile sensing) served as the means to obtain the optimal objective value (combined rotation and time score). These changes compared to the B-tests show the participants employed a trajectory modulation control policy. Moreover, they realised a reduced grip force in C-tests relative to B-tests is just enough for optimal task completion.

These findings suggest that our participants used trajectory modulation. Nonetheless, the impact of acceleration and its derivative on the object rotation is not like the reactive grip force control. There is a latency in the rotation with respect to the change in acceleration. We can speculate that humans employ a forward model for trajectory modulation, as opposed to reactive grip force control and/or simply memorising the movements. As such, they could efficiently reach the safety margin of grip force by predictive modulation of their hand motion profile. In the following, we explore our second research question hypothesising that this approach can be effective in robotic manipulation scenarios. We transferred this strategy for slip control to a robotic manipulation system and evaluated its performance in a range of manipulation tasks in the following. Inspired by the human study, our proposed controller learns to proactively make optimal adjustments to the robot hand’s trajectory instead of increasing the grip force for controlling rotational slip.

## 5. Discussion

Our results indicate that hand acceleration modulation plays a critical role in preventing slip during the pick-and-place task. The significant reduction in object rotation during the Controlled Tests supports the idea that humans use hand velocity modulation as a primary strategy for slip control when the grip force control is not effective. This finding challenges the conventional view that grip force is the only method employed by humans for slip prevention.

Interestingly, while grip force still plays a key role in object stabilization, participants in the Controlled Tests were able to minimize slip by modulating hand acceleration without significantly increasing grip force. This suggests that humans may employ a combination of control strategies to efficiently stabilise objects, balancing both grip force and motion adjustments.

Moreover, our findings suggest that trajectory modulation, in combination with predictive sensory feedback, is an effective strategy for controlling slip. This has important implications for robotics, where robotic systems have traditionally relied on reactive grip force control for slip prevention. The results of our human study provide a strong rationale for incorporating trajectory modulation into robotic slip control systems, particularly in environments requiring fine motor control and energy efficiency.

## 6. Conclusion

This study demonstrates that humans use hand acceleration modulation as an effective strategy for preventing slip during object manipulation tasks. Our results indicate humans may use combined hand movement modulation and grip force control to stabilize objects more efficiently than relying solely on grip force. These findings not only challenge the prevailing hypothesis in the field of slip control but also offer valuable insights for designing of robotic systems. By integrating trajectory modulation into robotic slip control policies, we can enhance robotic manipulation tasks, improving both stability and performance in dynamic environments.

Our research not only advances our understanding of human motor control strategies but also provides valuable insights for the design of more sophisticated and adaptive robotic manipulation systems. By demonstrating the importance of trajectory modulation in slip prevention, we offer a new approach for developing control algorithms that better mirror human manipulation strategies, potentially enhancing robotic performance in real-world applications.

Our results in [32] revealed hand movement modulations in predictive models can be integrated into robotic systems to optimize hand motion trajectories for slip prevention.

## Acknowledgments

This work was partially supported by the Centre for Doctoral Training, United Kingdom (CDT) in Agri-Food Robotics (AgriFoRwArdS) Grant reference: EP/S023917/1; Lincoln Agri-Robotics (LAR) funded by Research England; and by ARTEMIS project funded by Cancer Research UK C24524

/ A300038.

